# ATP Binding Cassette Proteins ABCG37 and ABCG33 are required for potassium-independent cesium uptake in Arabidopsis roots

**DOI:** 10.1101/823815

**Authors:** Mohammad Arif Ashraf, Sayaka Kumagai, Keita Ito, Ryohei Sugita, Keitaro Tanoi, Abidur Rahman

## Abstract

Radiocesium, accumulated in the soil by nuclear accidents is a major environmental concern. The transport process of cesium (Cs^+^) is tightly linked to the indispensable plant nutrient potassium (K^+^) as they both belong to the group I alkali metal with similar chemical properties. Most of the transporters that had been characterized to date as Cs^+^ transporters are directly or indirectly linked to K^+^. Using a combinatorial approach of physiology, genetics, cell biology and root uptake assay, here we identified two ATP-Binding Cassette (ABC) proteins, ABCG37 and ABCG33 as facilitators of Cs^+^ influx. The gain-of-function mutant of ABCG37 (*abcg37-1*) showed hypersensitive response to Cs^+^-induced root growth inhibition, while the double knock out mutant of ABCG33 and ABCG37 (*abcg33-1abcg37-2*) showed resistance. Single loss-of-function mutant of ABCG33 and ABCG37 did not show any alteration in Cs^+^ response. Short term uptake experiment with radioactive Cs^+^ revealed reduced Cs^+^ uptake in *abcg33-1abgc37-2* compared with wild type in presence or absence of K^+^. Potassium response and content were unaffected in the double mutant background confirming that Cs^+^ uptake by ABCG33 and ABCG37 is independent of K^+^. Collectively, this work identified two ABC proteins as new Cs^+^ influx carriers, which act redundantly and independent of K^+^ uptake pathway.

## Introduction

One of the most dangerous contaminants generated in a nuclear accident is cesium (Cs^+^), which shares similar chemical properties with potassium (K^+^). The stable isotope of cesium ^133^Cs is present at a very low concentration (less than 25 µg/g) in the soil and does not possess any major environmental concern (Cooghtrey et al., 1983). However, two radioisotopes of Cs (^134^Cs and ^137^Cs) are of environmental concerns due to their emission of harmful β and γ radiation, relatively long half-lives, and rapid incorporation into biological systems (White and Broadly, 2000, Kinoshita et al., 2011). In fact, consumption of agricultural product contaminated with radiocesium is the principal route of human exposure to this radionucleotide (Shaw and Bell, 1991), which has been shown to be related to increased risk of cancer. Trials of nuclear weapons, discharges from nuclear plants, and nuclear accidents such as the Chernobyl (Cooghtrey et al., 1983) and Fukushima incidents (Isaure et al., 2006) can result in large accumulation of radioactive Cs^+^ in the ground. This radioactive Cs^+^ is expected to be transmitted to the crops grown in the contaminated fields and cause serious health hazard for the population (Zhu and Smolders, 2000). For public health safety, the Japanese government restricted agricultural crop production on the soil containing more than 5000 Bq/kg (Yamaki et al., 2017). Hence, decontamination of the soil is one of the major priorities to keep the crop fields clean from radio cesium for producing crops free of contamination.

Radiocesium mostly accumulates on the surface of the soil (Fujiwara, 2013). Removal of top soil and transport to dedicated sites is one of the potential solutions for cleaning radioactive Cs^+^ contaminated field. Unfortunately, removal of soil from the surface, transportation and management is not an economic solution. Another technique is to use potassium fertilizer to reduce radiocesium toxicity (Zhu and Smolders, 2000). The classic work since 1940 suggest that Cs^+^ uptake is regulated through both K^+^ transporter and channels (Collander, 1941). Till today, most of the identified transporters that transport Cs^+^ are already reported K^+^ transporters. For instance, rice OsHAK1 (HIGH AFFINITY K^+^ TRANSPORTER 1) and OsHAK5 which are known K^+^ transporters can also function as cesium transporters (Nieves-Cordones et al., 2017; Rai et al., 2017). AtKUP/HAK/KT9 also functions as uptake carrier for potassium and cesium (Kobayashi et al., 2010). Hence, the symptoms of Cs^+^ intoxification can be reversed by supplying more K^+^ in the soil (Shaw and Bell, 1991). However, this strategy has major drawbacks; 1) the K^+^ transporter functioning at low external potassium concentration shows little discrimination against Cs^+^, while the K^+^ channel is dominant at high external K^+^ concentration with high discrimination against Cs^+^ (Zhu and Smolders, 2000), 2) high concentration of potassium itself is toxic to plants regardless of cesium concentration (Hampton et al., 2004), and 3) it is also economically impractical to provide large amount of potassium to soil. Further, it is a general consensus among the scientists that the mechanism by which Cs^+^ is taken up by plant roots is not completely understood. K^+^ transporters and channel are only partially responsible for Cs^+^ uptake and translocation (Zhu and Smolders, 2000). Taken together, these results suggest that there are alternative routes for Cs^+^ transport in plant which needs to be explored to understand the molecular mechanism of Cs^+^ uptake.

One of the major pathways for plant to detoxify toxic metals is through transporting to sequestering them in the vacuole. The ATP binding cassette (ABC) transporters, also called multidrug resistance proteins, are ubiquitous in plant and animal kingdom and play an important role in transporting various substances, including metals. For instance, tonoplast-localized AtABCC1 and AtABCC2 transport As (Song et al., 2010), Cd, and Hg (Park et al., 2012) inside vacuole. AtATM3/AtABCB25 is involved in cadmium transport (Kim et al., 2006). Plasma membrane-localized AtABCG36/AtPDR8 functions as efflux carrier of Cd (Kim et al., 2007). AtABCG40 has been shown to be linked to lead transport (Lee et al., 2005). These results suggest that diverse substrates, including multiple metals can be efficiently transported by the ABC superfamily transporters.

Although Cs^+^ has been suggested to compete with K^+^ for translocation, transcriptome analysis revealed that distinct set of genes show altered expression in Cs^+^ intoxicated plants compared with K^+^ starved plants (Hampton et al., 2004), confirming that Cs^+^ intoxication symptom in plants is not solely linked to K^+^. Interestingly, in Cs^+^ intoxicated plants, genes encoding ABC proteins showed altered expression pattern. At least 4 genes belong to this family showed 2.5-3 fold upregulation in response to Cs^+^ application but were not altered in K^+^ starvation (Hampton et al., 2004). In addition, a recent study using rice-transporter-enriched yeast expression library screening for Cs^+^ carrier identified one ABC transporter and one NRAMP transporter as functional Cs^+^ transporter in yeast (Yamaki et al., 2017). Taken together, these results suggest the possible existence of K^+^ independent transport system of Cs^+^ in plant and also identify ABC proteins as potential targets functioning as Cs^+^ transporters.

In the present work, we focused on ABC proteins to understand their roles in Cs^+^ transport. In plants, there are 8 subfamilies of ABC transporter proteins, namely ABCA-ABCH (Verrier et al., 2008). The *Arabidopsis* ABC superfamily comprises of 130 full molecular transporters containing at least two membrane spanning domains and two nucleotide binding folds. Among the ABC protein subfamilies, subfamilies B, C, and G have already been shown to possess metal transport activity (Lee et al., 2005; Kim et al., 2006; Kim et al., 2007; Song et al., 2010; Park et al., 2012). ABCC proteins are exclusively tonoplast localized and ABCB and ABCG are predominantly plasma membrane localized (Verrier et al., 2008). Since we are interested in Cs^+^ transport from soil, we focused on plasma membrane localized subfamilies (ABCB and ABCG). Through screening of 37 available ABCB and ABCG mutants root growth against a concentarion of Cs^+^ (1.5 mM) which inhibits 50 percent root growth in wild type, we identified ABCG33 and ABCG37 as potential Cs^+^ uptake carriers. The gain-of-function mutant of ABCG37 (*abcg37-1*) showed hypersensitive response to Cs^+^-induced root growth inhibition, while the double knock out mutant of ABCG33 and ABCG37 (*abcg33-1abcg37-2*) showed resistance. Single-loss-of function mutant of ABCG33 and ABCG37 (*abcg33-1*; *abcg37-2*) did not show any alteration in Cs^+^ response. Short term uptake experiment with radioactive Cs^+^ revealed reduced cesium uptake by *abcg33-1abgc37-2* compared with wild-type in presence or absence of K^+^. Potassium response and content were unaffected in the double mutant background confirming that Cs^+^ uptake by ABCG33 and ABCG37 is independent of K^+^. Collectively, this work identified two ABC proteins as new Cs^+^ uptake carriers, which act redundantly and independent of K^+^ uptake pathway.

## Results

### Cesium inhibits *Arabidopsis* primary root growth elongation

Metal stress or toxicity in plants is sensed via plant root and subsequently transported to other parts of the plant. To understand the effect of Cs^+^ on primary root growth, we performed both dose response and time course assay of CsCl using Arabidopsis wild-type (Col-0) seedlings. The time course and dose response data indicate that 1 mM and 1.5 mM cesium inhibits root growth approximately 50% after five days and three days incubation, respectively (Figure 1A). Because 1.5 mM Cs^+^ could inhibit ∼50% root growth in a shorter incubation period, we decided to use 1.5 mM cesium and three days incubation as an assay system for further experiments. This treatment affects the growth of the whole seedling including root and leaf. After 3 day incubation in 1.5 mM Cs^+^, root growth was inhibited and leaf chlorosis became apparent (Figures 1A and 1B).

**Figure 1.**
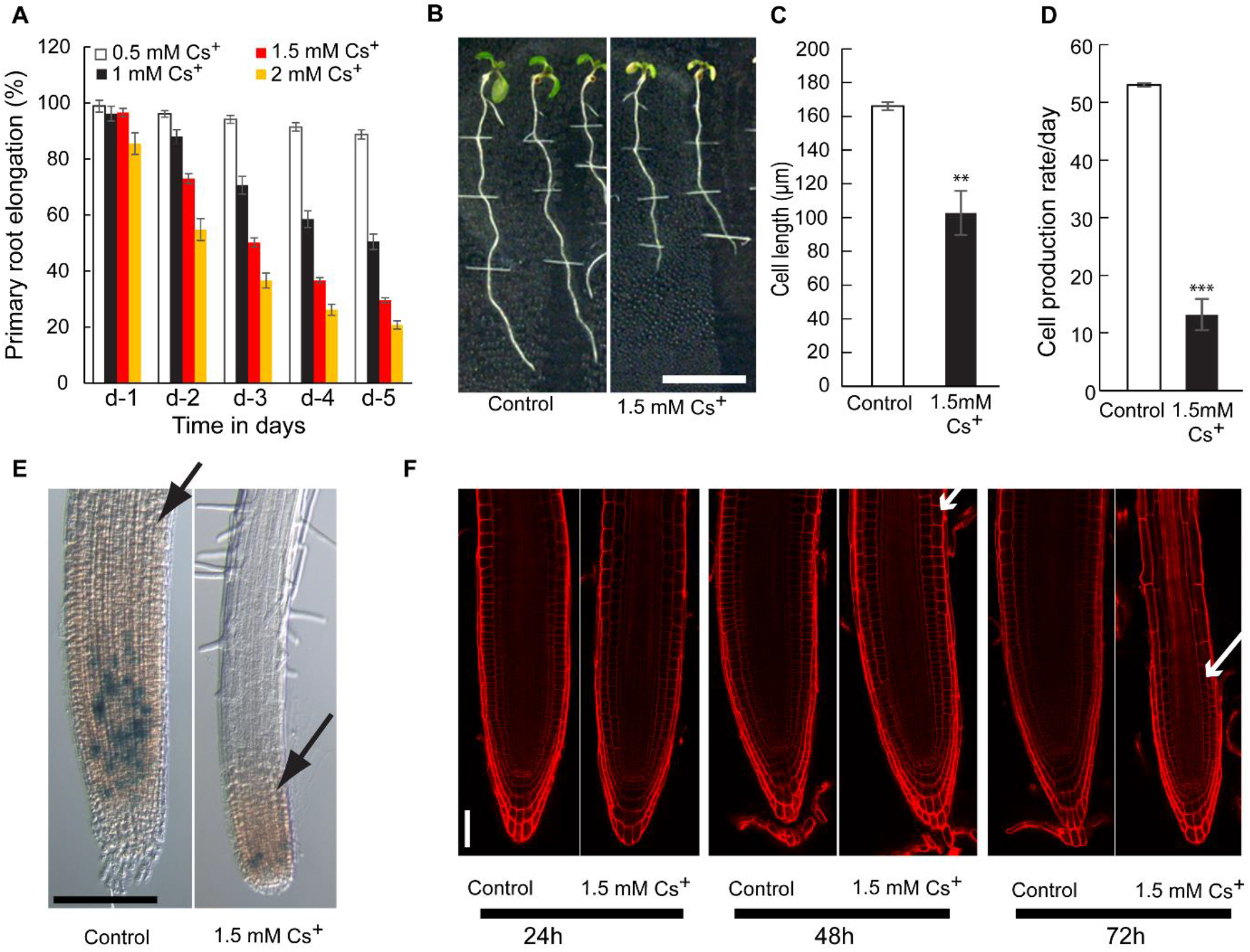
Effect of cesium on Arabidopsis root growth. **(A)** Primary root elongation of wild type (Col-0) in presence of Cs^+^. Three-day-old light grown wild-type (Col-0) seedlings were transferred to new agar plates with or without cesium (0.5, 1, 1.5 and 2 mM) and incubated at 23°C for five days. **(B)** Root phenotype of wild type (Col-0) after three days incubation in control and 1.5 mM Cs^+^ plates. **(C)** Cortical cell length of wild type (Col-0) after three days incubation in control and 1.5 mM Cs^+^ plates. **(D)** Cell production rate of wild type (Col-0) after three days incubation in control and 1.5 mM Cs^+^ plates. **(E)** Effect of cesium on expression of *CycB1;1-GUS* in the primary root tip. (F) Propidium idodide staining of wild type (Col-0). Three-day-old light grown wild type (Col-0) seedlings were transferred to new agar plates with or without 1.5 mM Cs^+^ and observed under confocal microscope at 24, 48 and 72h time point. Vertical bars mean ±SE from three independent experiments with 8 seedlings observed per experiment (**A, C,** D). Asterisks represent the statistical significance between treatments as judged by student’s *t*-test: **P < 0.01 and ***P < 0.001. Images are representative of at least three independent experiments with 5-8 seedlings observed per experiment (**B, E, F**). Meristem boundary is indicated by arrowheads. Bars= 10 mm for B and 100 µm for **E** and 50 µm for **F**.

The growth of primary root is a combination of cell division and elongation, which is further extended through differentiation (Rahman et al., 2007).Cesium-induced inhibition of primary root growth could be the consequence of either decreased cell number or shorter cell length or both. To answer this question, Cs^+^ treated seedlings were subjected to kinematic analysis (Rahman et al., 2007), which revealed that Cs^+^ inhibits both cell elongation and cell production (Figures 1C and 1D). The Cs^+^ effect on cell production rate was further confirmed by using G2-M phase cell cycle marker *CyclinB1;1::GUS* (Colón-Carmona *et al*., 1999), where only few cells were found to be mitotically active compared with the wild type (Figure 1E). The reduced cell division also affected the meristem growth as the reduction of meristem size was observed in Cs^+^ treated seedlings (Figure 1E and 1F). Although root growth was severely inhibited and root phenotype drastically changed after 3 day Cs^+^ incubation, the meristematic cells remain active as we did not observe any cell death by Propidium Iodide (PI) staining (Figure 1F).Taken together, these results suggest that cesium affects both cell division and elongation to inhibit the overall primary root elongation.

### Gain-of-function mutant *abcg37-1* is hypersensitive to cesium

In quest of finding new transporters for Cs^+^, we focused on ABC transporters as they have already been reported to transport metals such as As, Cd, Hg, Pb (Lee et al., 2005; Kim et al., 2006; Kim et al., 2007; Song et al., 2010; Park et al., 2012). The broader substrate specificity along with the altered expression pattern of some of the ABC family proteins under Cs^+^ intoxicated condition (Hampton et al., 2004) make them potential candidates for Cs^+^ transporter. To elucidate the roles of ABC proteins in Cs^+^ transport, we focused on plasma membrane localized subfamilies of ABC proteins ABCB and ABCG and screened available 20 ABCB and 17 ABCG mutants against 1.5 mM cesium for root growth response, which inhibits around 50% root growth in wild-type (Supplemental Figures 1A and 1B). The screening revealed four mutants (*abcb15-1*, *abcg36-1*, *abcg37-1*, *abcg42*) showing altered response to Cs^+^. Since we are interested in Cs^+^ uptake from soil, out of these four mutants, we selected *abcg37-1* (also known as *pdr9-1*) for further studies as it is a gain-of-function mutant (Ito and Gray, 2006), and showed hypersensitive response to Cs^+^. Time course and dose response analyses revealed that *abcg37-1* shows hypersensitive response to root growth at all concentrations and all incubation periods that we tested (Supplemental Figure 2). The Cs^+^ hypersensitivity in the gain of function mutant suggests that ABCG37 is possibly facilitating the uptake of Cs^+^, which results in increased root growth inhibition and leaf chlorosis (Figures 2B and 2C). If this is the case, then one would expect that the loss of function mutant will show opposite response. To validate this, we used several alleles of loss-of-function mutant of ABCG37 (*abcg37-2*, *abcg37-3*, *and abcg37-4*). These mutants show no visible root phenotype at control condition (Figure 2A). Unfortunately, loss-of-function mutants did not show cesium resistant phenotype.

**Figure 2.**
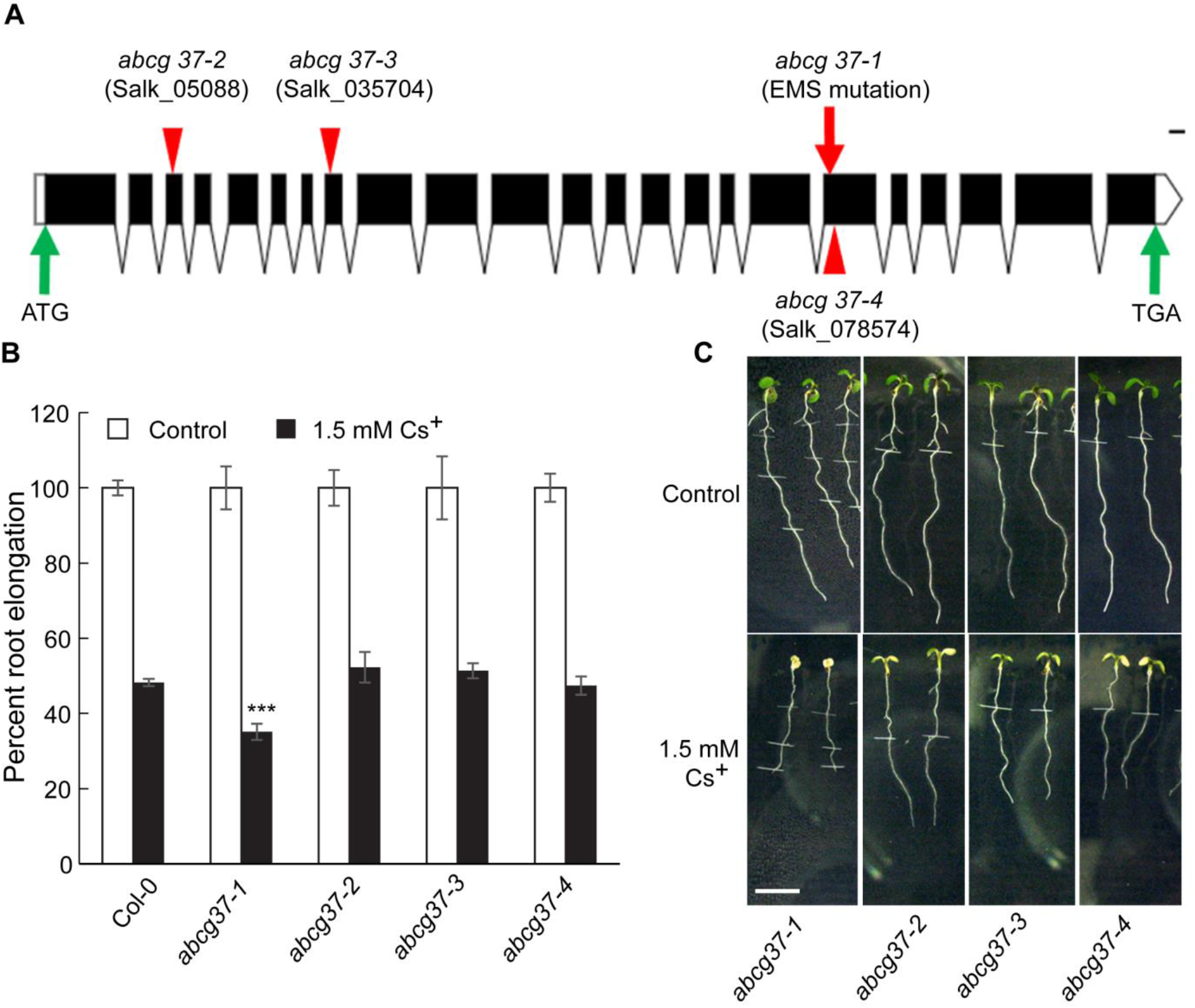
Gain-of-function mutant, *abcg37-1* shows hypersensitive response to cesium. (**A**) Representative diagram for *ABCG37* mutants used in this study. Solid black box and interrupted black line indicate exon and intron, respectively. (**B**) Primary root growth elongation of Col-0, *abcb37-1, abcg37-2, abcg37-3* and *abcg37-4* in presence of 1.5 mM Cs^+^. Three-day-old light grown seedlings were transferred to new agar plates with or without 1.5 mM Cs^+^ and incubated at 23°C for three days. Vertical bars mean ±SE from three independent experiments with 8 seedlings observed per experiment (**B**). Asterisks represent the statistical significance between treatments as judged by student’s *t*-test: ***P < 0.001 (**C**) Root phenotype of wild-type (Col-0) and *abcg37* mutants after three days incubation in control and 1.5 mM Cs^+^ plates. Control phenotype is demonstrated in Figure 1B. Images are representative of at least three independent experiments with 8 seedlings observed per experiment. Bar =10mm.

### ABCG37 and ABCG33 function redundantly as cesium uptake carriers

The unexpected wild-type like response of the loss-of-function ABCG37 mutants to cesium (Figure 2B) prompted us to hypothesize that loss of ABCG37 may be compensated by another ABC protein. For tonoplast localized arsenic transporter, *abcc1 abcc2* double mutant shows more severe phenotype compared to individual single mutants, suggesting the existence of redundant function of the ABC proteins (Song et al., 2010). To support the redundancy hypothesis, we searched the close homolog of ABCG37 based on phylogenetic analysis of 43 *Arabidopsis thaliana* ABCG proteins. ABCG33 turned out to be the closest homolog of ABCG37, residing in the same clade (Supplemental Figure 3A). They show ∼80% identity at protein level (Supplemental Figure 3B).

Double knockout homozygous *abcg33-1 abcg37-2* mutant was generated to test the redundant functional hypothesis. Before crossing, using gene expression analysis we confirmed that both *abcg33-1* and *abcg37-2* mutants are null mutants (Supplemental Figure 4). Consistent with our hypothesis, we found that *abcg33-1 abcg37-2-*11, *abcg33-1 abcg37-2-*21 and *abcg33-1 abcg37-2-23* (three independent double mutant lines obtained from three independent crossing between *abcg37-2* and *abcg33-1*) showed resistance to cesium for both root growth inhibition and leaf chlorosis (Figures 3A and 3B). The time course and dose response assay revealed that the double mutants showed strong resistance to lower concentration of Cs^+^-induced root growth inhibition at all incubation period (Supplemental Figures 5A and 5B). The resistant phenotype becomes weaker with progressively higher concentration of Cs^+^ (Supplemental Figure 5C). However, at early incubation time point (like 2 days incubation in Cs^+^), the double mutants showed strong resistance to Cs^+^ irrespective of the concentration (Supplemental Figure 5). ABCG37 has been shown to function as IBA efflux carrier (Růžička et al., 2010). Since majority of IBA functions in plant development through conversion to IAA (Růžička *et al*., 2010; Frick and Strader, 2017), we investigated whether *abcg37*, *abcg33* and *abcg33 abcg37* mutants show altered root growth response to exogenous IAA. All these mutants show wild-type like response to IAA-induced root growth inhibition, indicating that the altered response of these mutants to Cs^+^ is not linked to the major endogenous form of auxin, IAA (Supplemental Figure 6). Collectively, these results confirm that ABCG33 and ABC37 act redundantly to uptake Cs^+^.

**Figure 3.**
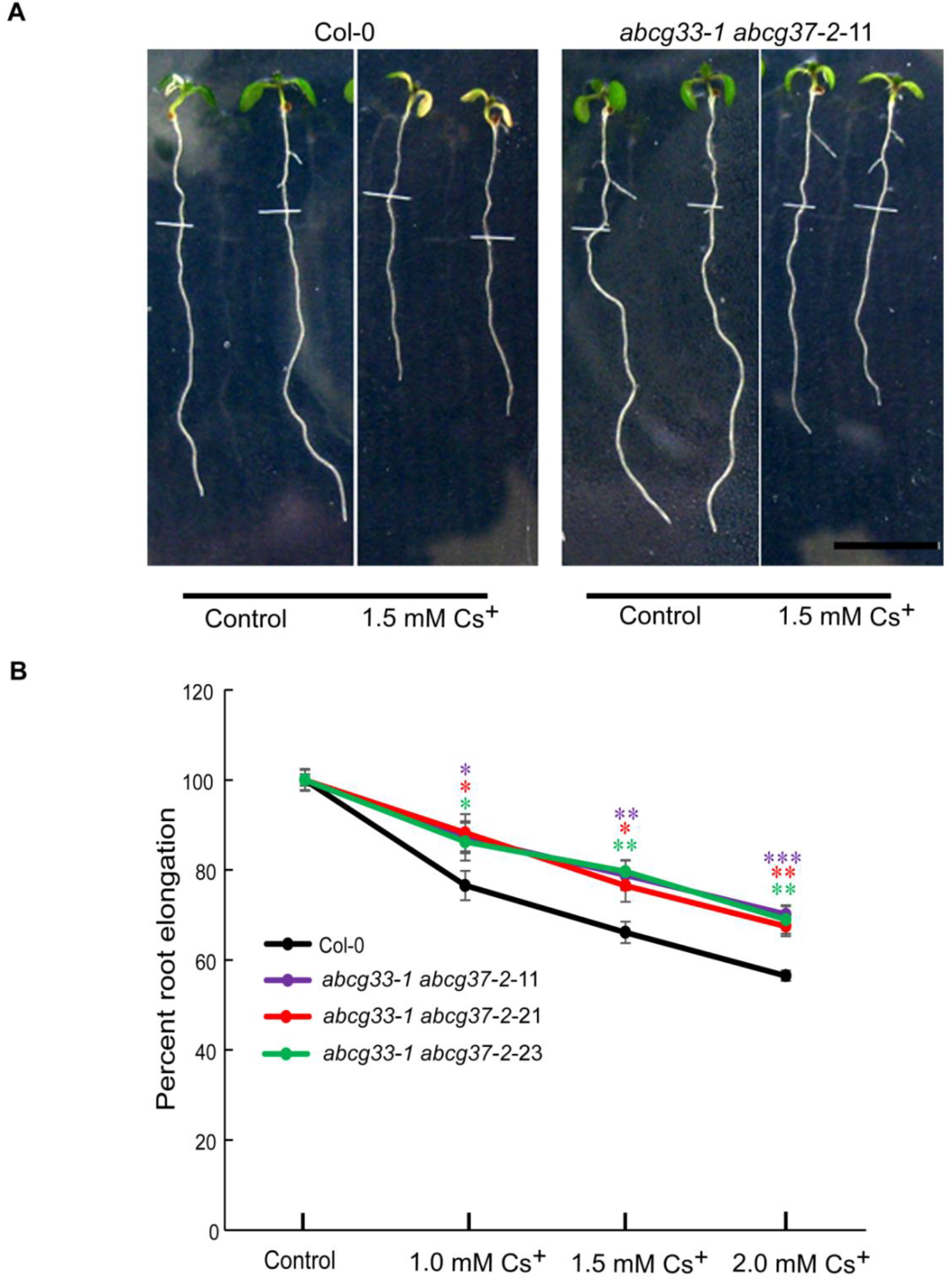
ABCG37 and ABCG33 work redundantly as cesium transporter. (**A**) Root phenotype of wild type (Col-0) and *abcg37-2 abcg33-1* −11 after three days incubation in control and 1.5 mM Cs^+^ plates. Images are representative of at least three independent experiments with 8 seedlings observed per experiment. Bar =10 mm. (**B**) Primary root elongation of Col-0, *abcb37-2 abcg33-1*-11, *abcb37-2 abcg33-1*-21 and *abcb37-2 abcg33-1-*23 in presence of 1, 1.5 and 2 mM Cs^+^. Three-day-old light grown seedlings were transferred to new agar plates and incubated at 23°C for two days. Vertical bars mean ±SE from three independent experiments with 8 seedlings observed per experiment. Asterisks represent the statistical significance between treatments as judged by student’s *t*-test: *P < 0.05, **P < 0.01 and ***P < 0.001.

### Cesium regulates the expression of ABCG37 and ABCG33 proteins at translational level

Many of the metals (Cd^2+^, Pb^2+^) regulate the gene expression of their corresponding ABC transporters (*AtABCG36*, *AtABCB25*, and *AtABCG40*) (Lee et al., 2005; Kim et al., 2006; Kim et al., 2007). Furthermore, differential expression of a group of Arabidopsis genes, including ABC proteins was observed in cesium intoxicated plants (Hampton et al., 2004). To test whether cesium regulates the expression of *ABCG37* and *ABCG33*, we used quantitative real time PCR. However, compared with wild-type, we did not observe any significant changes in the transcripts of *ABCG37* and *ABCG33* in Cs^+^ treated plants (Supplemental Figure 7). This observation is not inconsistent as in few cases it has been shown that transporters transcriptions are not affected by the application of the substrate. For instance, arsenite has no effect on expression of *AtABCC1* and *AtABCC2* (Song et al., 2010); rice cesium transporters are not transcriptionally altered by cesium (Yamaki et al., 2017).

The quantitative real-time PCR analyses of *ABCG37* and *ABCG33* indicated that ABCG37 and ABCG33 are not under direct transcriptional regulation of cesium. To understand whether Cs^+^ regulates ABCG33 and ABCG37 at translational level, we monitored the cellular expression of these proteins using GFP tagged lines. Co-localization study with PI confirmed that both ABCG33 and ABCG37 are plasma membrane localized protein (Supplemental Figure 8). Interestingly, we found that both ABCG37-GFP and ABCG33-GFP intracellular signals are severely reduced by 1.5 mM cesium treatment in a time dependent manner. The reduction of the GFP signal is maximum on day three but the Cs^+^ effect on these proteins are apparent from day 2 (Figures 4A and 4B). To clarify the specificity of Cs^+^ effect on ABCG33 and ABCG37 proteins, we used two root specific membrane proteins, PIN2-GFP (Xu and Scheres, 2005) and EGFP-LTI6b (Kurup et al., 2005). Cs^+^ slightly reduced PIN2-GFP signal but did not affect the LTI6b expression (Figure 4B; Supplemental Figure 9). These results nullify the possibility that the Cs^+^-induced inhibition of ABCG33 and ABCG37 expression is due to toxicity, rather it suggests that Cs^+^ selectively inhibits a subset of membrane proteins.

**Figure 4.**
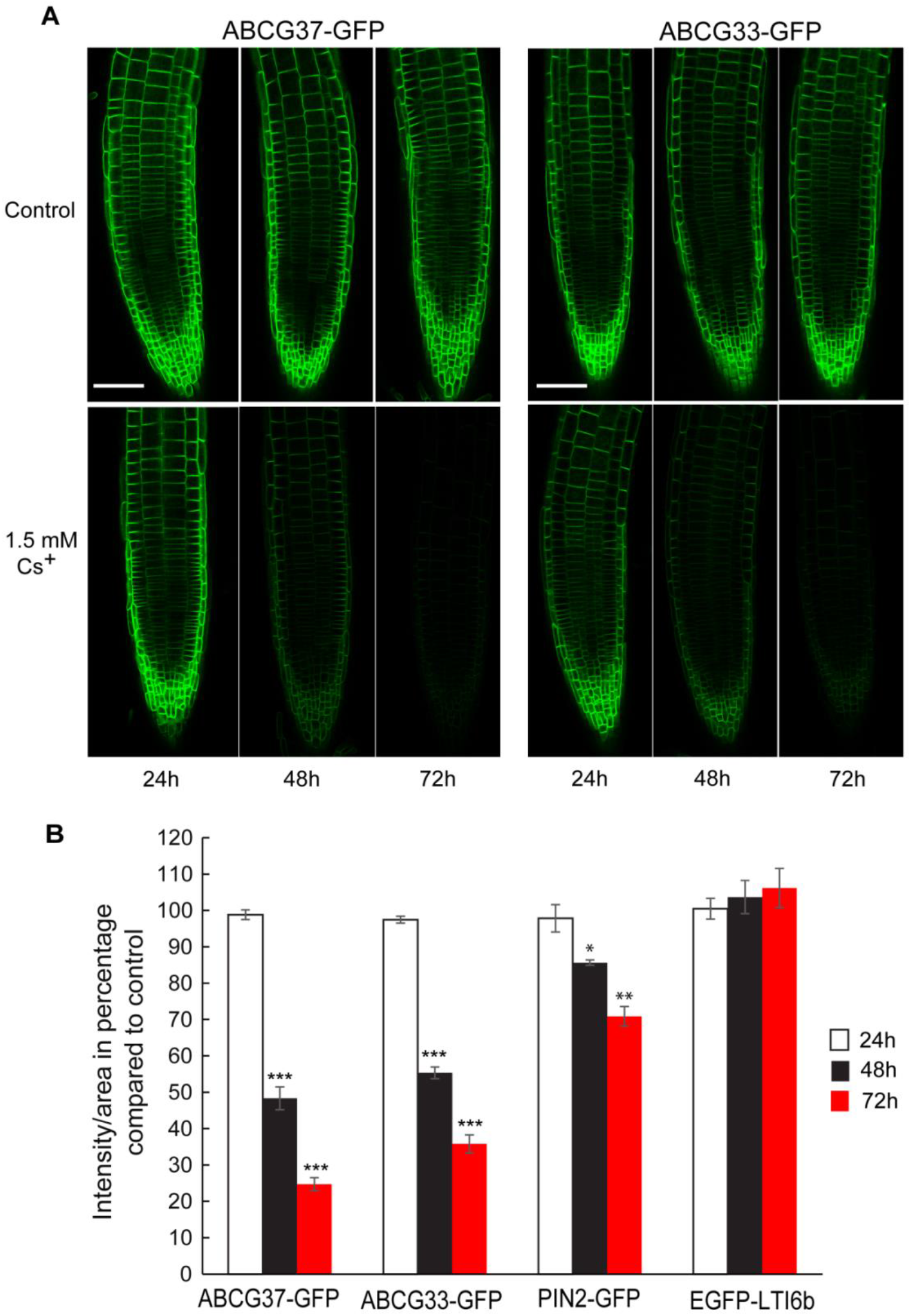
Cesium inhibits the expression of ABCG33 and ABCG37. Three-day-old light grown ABCG37-GFP and ABCG33-GFP seedlings were transferred to control and 1.5 mM Cs^+^ containing plates. (**A**) GFP fluorescence of ABCG37-GFP and ABCG33-GFP were observed at 24, 48 and 72h time point. The images are single confocal sections representative of three independent experiments with 5 seedlings observed per treatment for each experiment. Bars=50µm. (**B**) Quantification of GFP signal in ABCG37-GFP, ABCG33-GFP, PIN2-GFP and EGFP-LTI6b from control and 1.5 mM Cs^+^ containing plates at 24, 48 and 72h time point. Quantification was performed from three independent experiments with 5 seedlings observed per treatment for each experiment. Vertical bars mean ±SE. Asterisks represent the statistical significance between treatments as judged by student’s *t*-test: *P < 0.05 and ***P < 0.001.

### ABCG37 and ABCG33 are functionally redundant cesium influx carriers

The physiological, molecular and cellular analyses identified ABCG33 and ABCG37 as potential Cs^+^ influx carriers. However, evidence for direct influx activity remains lacking. The uptake activity of a protein can be assessed by either using heterologous expression system or direct *in planta* uptake assay. To confirm the potential role of ABCG33 and ABCG37 in Cs^+^ uptake, we first performed short term uptake assay in Arabidopsis roots using radio labeled cesium (^137^Cs).

Cs^+^ uptake capacity of wild type and double mutants was measured by direct uptake assay using radioactive ^137^Cs^+^. The Cs^+^ uptake was observed for 2h in presence or absence of K^+^ at various Cs^+^ concentrations ranging from low to high (Figure 5). The short-term assay system eliminates the possible non-specific effects on transport. Irrespective of the presence or absence of K^+^, *abcg33-1 abcg37-2* double mutants showed a reduced cesium uptake capacity compared with wild-type at all cesium concentrations we tested (Figure 5A, 5B, 5C). To confirm that the double mutant lines truly lack the Cs^+^ uptake capacity, we compared Cs^+^ content in roots of the wild type and *abcg33-1 abcg37-2* mutants after 3 days Cs^+^ incubation using ICP-MS. Compared with wild type, Cs^+^ content was significantly reduced in *abcg33-1 abcg37-2* mutant lines at all K^+^ concentration we tested (Figure 5D). The short-term uptake data along with ICP-MS data suggest that ABCG37 and ABCG33 function as cesium influx carriers.

**Figure 5.**
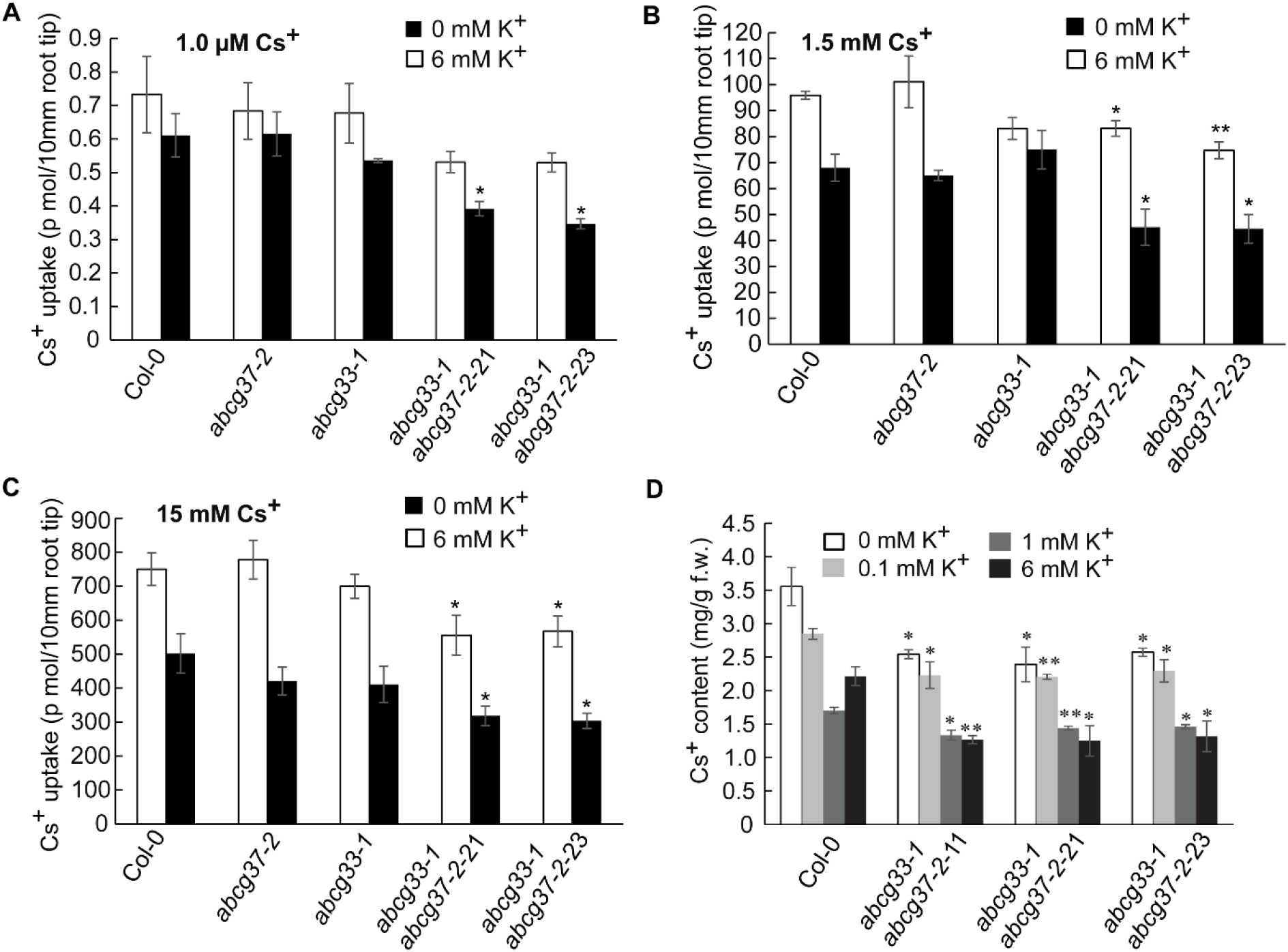
ABCG33 and ABCG37 function as cesium influx carriers. (**A-C**) Short term uptake assay of ^137^Cs^+^ in Col-0, *abcg37-2 abcg33-1* (11), *abcg37-2 abcg33-1* (21) and *abcg37-2 abcg33-1* (23) root tip. Four-day-old light grown Col-0, single and double mutant seedlings were incubated for 2h at 10μM, 1.5mM,15 mM ^137^Cs^+^ in presence (6mM) or absence of K^+^. After the incubation, seedlings were washed three times and 10mm root tip from 10 roots were excised for each sample and radioactivity was counted using a scintillation counter. Data are the averages from at least 3 independent experiments, and expressed as per mm root tip. Vertical bars mean ±SE. Asterisks represent the statistical significance between genotypes as judged by student’s *t*-test: *P < 0.05, **P < 0.01 (**D**) Cesium content in Col-0 and *abcg* double mutants. Three-day-old light grown Col-0 and double mutant seedlings were transferred to 1.5mM Cs^+^ plates in presence of various K^+^ concentrations, and incubated for three days. Cs^+^ content of whole seedling was measured by ICP-MS. The data were obtained from three independent experiments. Vertical bars mean ±SE. Asterisks represent the statistical significance between treatments as judged by student’s *t*-test: *P < 0.05, and **P < 0.01.

### ABCG37 does not show Cs^+^ uptake activity in *Saccharomyces cerevisiae*

To evaluate ABCG37 activity in the Cs^+^ uptake in a heterologous expression system, we expressed ABCG37 under GAL1 promoter in INVSc1 strain of budding yeast *Saccharomyces cerevisiae*. Unfortunately, no significant differences were observed for Cs^+^ uptake between ABCG37 expressing cells and control cells at all Cs^+^ concentrations we tested (Supplemental figure 10). These results suggest that ABCG37 needs other components to function as CS^+^ influx carrier, which is present in plant cell but absent in yeast system. This finding is not surprising as it has already been demonstrated that several plant proteins fail to transport corresponding metals when expressed in a heterologous system (Gaymard et al. 1996; Formentin et al. 2004; Ma et al. 2008), or transport with a limited capacity only after co-expressing with other proteins (Song et al. 2010; Park et al. 2012). Expression of ABCG37 in *Saccharomyces cerevisiae* resulted in mis localization of the protein to endoplasmic reticulum instead of plasma membrane and did not show any efflux activity for IBA (Růžička et al. 2010). When ABCG37 was expressed in *Schizosaccharomyces pombe*, the plasma membrane localization of ABCG37 could be achieved but the efflux activity was observed only at unusually high concentration (250 μM) of IBA (Růžička et al. 2010), although ABCG37 showed IBA efflux activity at much lower concentration in planta. ABCG33 expression in *Saccharomyces cerevisiae* was unsuccessful. Taken together, these results suggest that investigating the transporter activity of a plant protein in heterologous system is tricky and may not always mimic *in planta* results.

### Potassium response is unaltered in *abcg33abcg37* double mutant

Because of the structural similarity of Cs^+^ with K^+^, most of the Cs^+^ transporters found today also transport K^+^. Here we identified two ABCG proteins that modulate Cs^+^ influx in Arabidopsis roots. To verify that these proteins solely uptake Cs^+^ but not K^+^, we measured the endogenous K^+^ content in wild-type and double mutants using ICP-MS and also compared the root growth response to a concentration of K^+^ that inhibits 50% root growth in wild type. ICP-MS data revealed no significant difference in K^+^ content between the wild type and double mutants grown at different concentrations of K^+^ (Figure 6A). Consistently, the double mutants respond to exogenous K^+^ for root growth like wild-type (Figure 6B). These results provide strong evidence in support of the hypothesis that ABCG33 and ABCG37 are Cs^+^ specific uptake carriers and act independent of K^+^.

**Figure 6.**
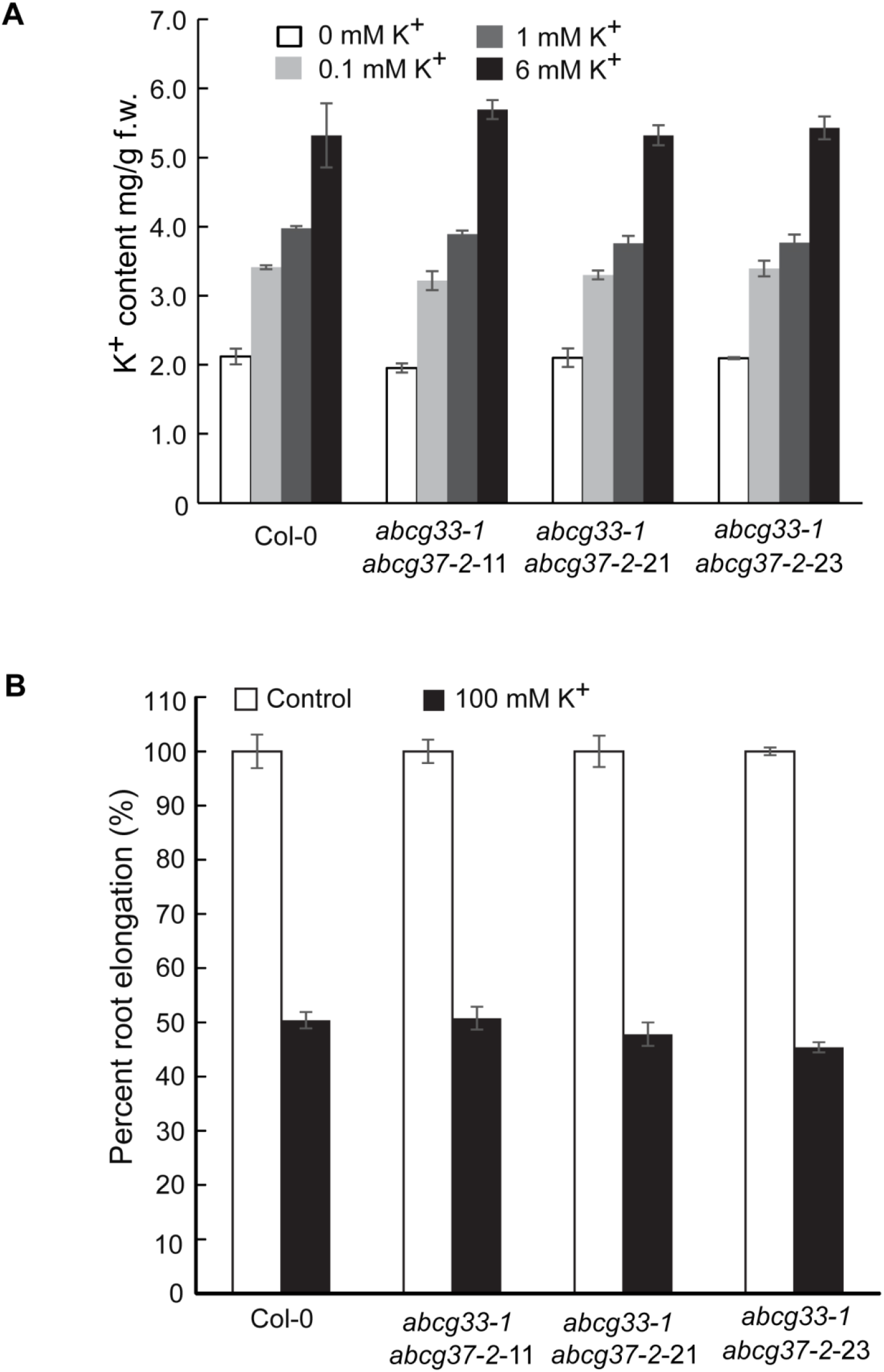
ABCG33 and ABCG37 are potassium-independent cesium transporters. **(A)** Potassium content in Col-0, *abcg37-1*, *abcg* double mutant seedlings. Three-day-old light grown Col-0 and double mutant seedlings were transferred to fresh plates containing various concentrations of K^+^, and incubated for three days. K^+^ content of whole seedling was measured by ICP-MS. (**B**) Primary root elongation response of Col-0, *abcg33-1 abcg37-2*-11, *abcg33-1 abcg37-2*-21 and *abcg33-1 abcg37-2*-23 in presence of 100mM potassium chloride. Four-day-old light grown Col-0 and double mutant seedlings were transferred to 100 mM potassium chloride containing plates and incubated for three days. For both (**A**) and (**B**) vertical bars mean ±SE from three independent experiments. The K^+^ content and percent root elongation in double mutant are statistically non-significant compared with wild-type as judged by the student’s *t*-test.

## Discussion

Cesium belongs to group I alkali metal with chemical properties similar to K^+^. Not surprisingly, the transport and translocation of Cs^+^ in plant is tightly linked to K^+^. Most of the transporters that had been characterized to date as Cs^+^ transporters are directly or indirectly linked to K^+^. Recently identified cesium transporters from Arabidopsis, rice, tomato and other system are mostly well studied potassium transporters, such as *OsHAK1* (Nieves-Cordones et al., 2017; Rai et al., 2017), *AtKUP/HAK/KT9* (Kobayashi et al., 2010), *OsHAK5*, *OsHAK17* (Yamaki et al., 2017), *SIHAK5* (Ródenas et al., 2017). Even the ecotype screening for cesium transporters pointed out *AtCNGC1* (*CYCLIC-NUCLEOTIDE-GATED CHANNEL 1*) as potential causal gene, which is the close homolog of *AtCNGC2*, a known K^+^ transporter (Kanter et al., 2010). In another study, *OsSOS2* had been shown to transport Cs^+^ (Ishikawa et al., 2017). However, *OsSOS2* was found to be indirectly linked to K^+^ as in *OsSOS2* mutant, K^+^ and Na^+^ transporter genes (*OsHAK1*, *OsHAK5*, *OsAKT1*, *OsHKT2;1*) are downregulated at low K^+^/Na^+^ (Ishikawa et al., 2017). It is predicted that Cs^+^ uptake is reduced due to the lower expression of these transporters in presence of the K^+^/Na^+^ imbalance (Ishikawa et al., 2017). Altogether, these results suggest a possible single pathway of Cs^+^ transport in plants. However, this idea is partially correct as all the recently identified low cesium concentration mutants function only under low K^+^ content (Qi et al., 2008; Nieves-Cordones et al., 2017, Ishikawa et al., 2017; Rai et al. 2017). At higher K^+^ concentrations, the selectivity for K^+^ strongly increases and this selective discrimination indicates the existence of alternate route of Cs^+^ transport in planta.

In the present study, we identified two ABC proteins (ABCG33 and ABCG37) that function as cesium influx carriers. The broader specificity of ABC proteins make them excellent transporter of many metals, hormones and chemical compounds. Previously, ABCG37 has been reported to act as efflux carrier of IBA (Růžička et al., 2010). Here we demonstrated that ABCG37 and ABCG33 function as cesium influx carriers. This observation is not inconsistent as the same ABC transporter can transport various substrates. For instance, ABCC1 and ABCC2 transport arsenic, mercury, and cadmium (Lee et al., 2005; Kim et al., 2006; Kim et al., 2007; Song et al., 2010; Park et al., 2012). AtPDR8/ABCG36 transports cadmium, lead and IBA (Kim et al., 2007, Růžička et al., 2010). AtPDR12/ABCG40 transports both cadmium and ABA (Lee et al., 2005; Kang et al., 2010) Similarly, ABCG37 also shows a wide substrate specificity and transports IBA, 2,4-D and coumarin (Ito and Gray, 2006; Růžička et al., 2010; Fourcroy et al., 2014). These substrates are not even distantly related to each other structurally. In contrast, the functional role of ABCG33 is yet undefined although it may function in transporting monolignol (Schuetz et al., 2014). The findings that ABCG37 can function as both influx (for Cs^+^) or efflux (for 2,4-D and IBA) is not inconsistent as based on the substrate, ABC transporters may act as influx, efflux or both at the same time. PGP4/ABCB4 was demonstrated as influx (Terasaka et al., 2005), efflux (Cho et al., 2007) and both influx/efflux (Yang and Murphy, 2009; Kubeš et al., 2012; Swarup and Péret, 2012) carriers of auxin. Similarly, PDR12/ABCG40 has been shown to function as efflux carrier for cadmium but as influx carrier for ABA (Lee et al., 2005; Kang et al., 2010)

Since ABCG37 has been shown to be linked to transport 2,4-D and IBA (Ito & Gray, 2006; Růžička *et al*., 2010), it may be hypothesized that the Cs^+^-induced inhibition of root growth is linked to auxin. Several lines of evidence argue against this notion; 1) The phenotype of Cs^+^-induced inhibition of plant growth is completely different than the auxin-induced inhibition. Long term Cs^+^ treatment results in leaf chlorosis, which is never induced by auxin treatment (Figures 1,2,3). Additionally, the double mutant *abcg33-1abcg37-2* shows strong resistance to Cs^+^-induced leaf chlorosis (Figure 3). 2) Auxin-induced root growth inhibition is associated with an increase in lateral root number. In case of Cs^+^-induced inhibition, the lateral root development is suppressed (Figures 1,2,3), indicating that these two chemicals use distinct pathways to inhibit the root growth. 3) All the single or double mutants of ABCG33 and ABCG37 show wild-type like root elongation response to major endogenous auxin IAA (Supplemental Figure 6). Previously it was demonstrated that ABCG37 mutants show wild-type like respons to IAA (Ito & Gray, 2006; Růžička *et al*., 2010), which is consistent with our results. Further, 2,4-D is not an endogenous auxin and hence Cs^+^-induced inhibition is not linked to 2,4-D. In case of IBA, it is believed that majority of IBA functions in plant development through conversion to IAA (Růžička *et al*., 2010; Frick and Strader, 2017). Since all the ABCG33 and ABCG37 single and double mutants respond to IAA like wild-type, it is reasonable to speculate that the response of these mutants to Cs^+^ is independent of auxin.

Our physiological, genetic, cell biological and direct transport assay data indicated that ABCG33 and ABCG37 function redundantly in uptaking Cs^+^. Additionally, the reduction in Cs^+^ uptake in *abcg33 abcg37* double mutant is independent of K^+^ availability in the media as similar reduction in Cs^+^ uptake was observed in presence or absence of K^+^ (Figure 5).The specificity of these proteins to uptake only Cs^+^ but not K^+^ was also confirmed by measuring K^+^ concentration and root growth response to exogenous K^+^ in the double mutant lines (Figure 6). Unfortunately, ABCG37 did not show any Cs^+^ influx activity in heterologous system. This is not inconsistent as the expression of plant proteins in heterologous system widely varies depending on the system that is used, and in many cases plant proteins do not express in heterologous system. For instance, AKT1 and DKT1 showed disturbed cell membrane electrical stability and inactivity when expressed in oocyte system (Gaymard et al., 1996; Formentin et al., 2004). Dreyer et al. also elegantly showed the problems with the heterologous expression system in expressing plant potassium transporters highlighting the differences between the expression systems (Dreyer et al., 1999). Lsi2, which functions as arsenite and silicon transporter in planta, did not show any transport activity when it was expressed using heterologous systems (Ma et al., 2008). Bernaudat et al. performed extensive analyses of expression of plant membrane proteins in wide range of heterologous expression systems and clearly showed that the expression varies depending on the expression system and in some cases the plant proteins simply do not express in heterologous system. The authors concluded that “with heterologous protein expression, the recombinant produced does not always truly resemble the native protein. Conditions that produce the largest amount of protein do not necessarily generate functional proteins” (Bernaudat et al., 2011). This conclusion was also supported by several published work for mammalian transporter proteins (Lenoir et al., 2002; Griffith et al., 2003; Tate et al., 2003; Bonander et al., 2005; Midgett and Madden, 2007). Additionally, since the activity of the transporters in plants may be modulated by their interactions with other proteins, the inability of the transporters to transport their respective substrates when expressed in heterologous system is not unusual as the interacting proteins that are required for proper functioning may be missing in heterologous expression systems (Zourelidou et al., 2014). Consistently, ABCC1 and ABCC2 could transport arsenic only when they were expressed in phytochelatins (PCS) producing yeast strain (Song et al., 2010). Similar results were observed for mercury. ABCC1 and ABCC2 transported more mercury when it was expressed in PCS producing yeast compared with yeast strain that does not produce any PCS (Park et al., 2012). Interestingly, the same ABCC1 and ABCC2 failed to transport any cadmium even when they were expressed in PCS producing yeast strain, although they could transport cadmium in planta (Perk et al., 2012). Hence, the inability of the plant transport proteins to transport substrate when expressed in heterologous system does not exclude them to function as transporters *in planta*.

Previously, plasma membrane-localized metal transporters were established based on only *in planta* experimental system such as using radioisotope for transport assay and measuring content through ICP-MS. For instance, AtPDR8/ABCG36 and AtATM3/ABCG25 were identified as Cd transporter based on radioactive ^109^Cd transport assay and ICP-MS (Kim et al., 2006; Kim et al., 2007). Additionally, AtPDR12/ABCG40 was identified as Cd transporter based on ICP-MS analysis, where increased or decreased content of Cd was observed for *atpdr12-1* and overexpression line of AptPDR12, respectively (Lee et al., 2005). These results along with the observed in planta Cs^+^ influx activity of ABCG33 and ABC37 characterized these two ABC proteins as potential candidates for Arabidopsis specific Cs^+^ influx carriers.

Our finding that ABC proteins may function as Cs^+^ specific influx carriers is supported by the recent identification of cesium transporters from the screening of rice-transporter-enriched yeast expression library where rice OsABCG45 has been shown to be a functionally active cesium transporter in yeast (Yamaki et al., 2017). OsABCG45 shows ∼50% homology with both ABCG33 and ABCG37 from Arabidopsis. Cs^+^ intoxification results in upregulation of *AtABCG16*, *AtABCA7* and *AtNAP5* in Arabidopsis (Hampton et al., 2004). In addition, during our screening, we found additional three loss of function ABC mutants; *abcb15, abcg36* and *abcg42* showing hypersensitive response to Cs^+^-induced inhibition of root growth. Collectively, these results suggest that beside ABCG33 and ABCG37, other ABC transporters may function as cesium transporters.

Another possibility remains open that the reduced uptake or accumulation of Cs^+^ in the *abcg33abcg37* double mutant may be due to nonspecific effect of the mutations to other K^+^ transporters. Although we do not completely rule out the possibility, several lines of experimental evidence go against this possibility. Measurement of K^+^ content in the double mutant in presence of low to high concentrations of K^+^ did not reveal any alteration in the potassium accumulation in the double mutant compared with the wild-type (Figure 6A). The potassium content was measured after 3-day treatment confirming that there are no non-specific changes in other transport proteins that may involve in transporting K^+^. Further, the root growth response of double mutant under K^+^ showed no changes in the root elongation response suggesting that these mutations do not alter potassium response (Figure 6B). Measurement of Cs^+^ content in presence of variable concentrations of K^+^ revealed a similar trend of decreasing Cs^+^ content under high K^+^ condition both in wild type and double mutant (Figure 5D). If there was a change in K^+^ transport in double mutant, one would expect to observe a clear difference in double mutant. Based on these multiple lines of evidence, we concluded that the double mutant has an unaltered K^+^ transport.

Although it has been shown that both low and high Cs^+^ can alter the gene expression in Arabidopsis (Hampton et al., 2004; Sahr et al., 2005a, Sahr et al., 2005b), ABCG33 and ABCG37 are not under the transcriptional regulation of Cs^+^. Cesium alters the expression of ABCG33 and ABCG37 protein in a time dependent manner. Since ABCG33 and ABCG37 function as potential Cs^+^ influx carriers, one plausible explanation is plants shut off the transporters to protect it from long term toxicity. This process is possibly regulated by the cesium-mediated post-translational regulation of transporters. However, the mechanism of this regulation is still unclear. Similar observation was reported for BRASSINOSTEROID INSENSITIVE 1(BRI1) receptor regulation at prolonged ambient temperature. Although 26°C incubation did not alter the transcript of *BRI1*, the protein expression was shown to be decreased in a time dependent manner (Martins et al., 2017). Polyubiquitin-dependent endocytosis and degradation have been suggested to regulate this process (Martins et al., 2017). Cesium toxicity has already been reported to induce proteolytic degradation of AGO1 possibly through autophagy (Jung et al., 2015). The ferrous Fe uptake transporter IRON-REGULATED TRANSPORTER1 (IRT1) is shown to be rapidly and constitutively degraded through a ring E3 ubiquitin ligase IRT Degradation Factor 1 (IDF1)-mediated pathway to maintain cellular Fe homeostasis (Shin et al., 2013). IRT1 also serves as a transceptor, directly sensing non-iron metals using a histidine-stretch in IRT1, and regulates its own degradation by differential ubiquitination upon metal stress to maximize iron uptake while limiting the absorption of highly reactive and potentially toxic non-iron metals. (Dubeaux et al, 2018). Recent studies on ethylene signaling provided further insight about translational regulation. C-terminal end of EIN2 (EIN2-CEND) interacts with 3’UTR of EBF1/2 and represses the translation of EBF1/2 by directing them towards P body for RNA decay. As a result, EBF1/2 become unable to degrade EIN3/EIL1, major transcription factors for ethylene response (Li et al., 2015; Merchante et al., 2015). These results indicate the existence of various mechanisms for post translational regulation of proteins. Future research focusing on understanding the cesium mediated translational regulation will shed light on intracellular Cs^+^ signaling events.

The present work identified the long-sought K^+^ independent uptake carriers of Cs^+^. ABCG protein mediated Cs^+^ uptake facilitates the Cs^+^ transport along with known K^+^ transporters. Based on the results, we developed a working model where cesium uptake and transport inside the cell are mediated by potassium transporters, ABC transporters (ABCG37 and ABCG33) and other unknown transporters (Figure 7). Hence, in the double knockout mutant we did not observe complete resistance to cesium. Bioremediation of the contaminated soil is a clever and affordable technology, which is largely unused for Cs^+^ removal because of the similarity in chemical properties of Cs^+^ to K^+^, a major macronutrient for plant growth. Removal of K^+^ transporters affects the plant growth and also low K^+^ content soil is also a common requirement to transport contaminated Cs^+^ (Nieves-Cordones et al., 2017; Rai et al., 2017; Ródenas et al., 2017; Yamaki et al., 2017). Future research aiming in generating new transgenic plants manipulating both K^+^ transport pathway and the newly identified ABC protein mediated Cs^+^ transport pathway will facilitate the efficient removal of Cs^+^ from the contaminated soil.

**Figure 7.**
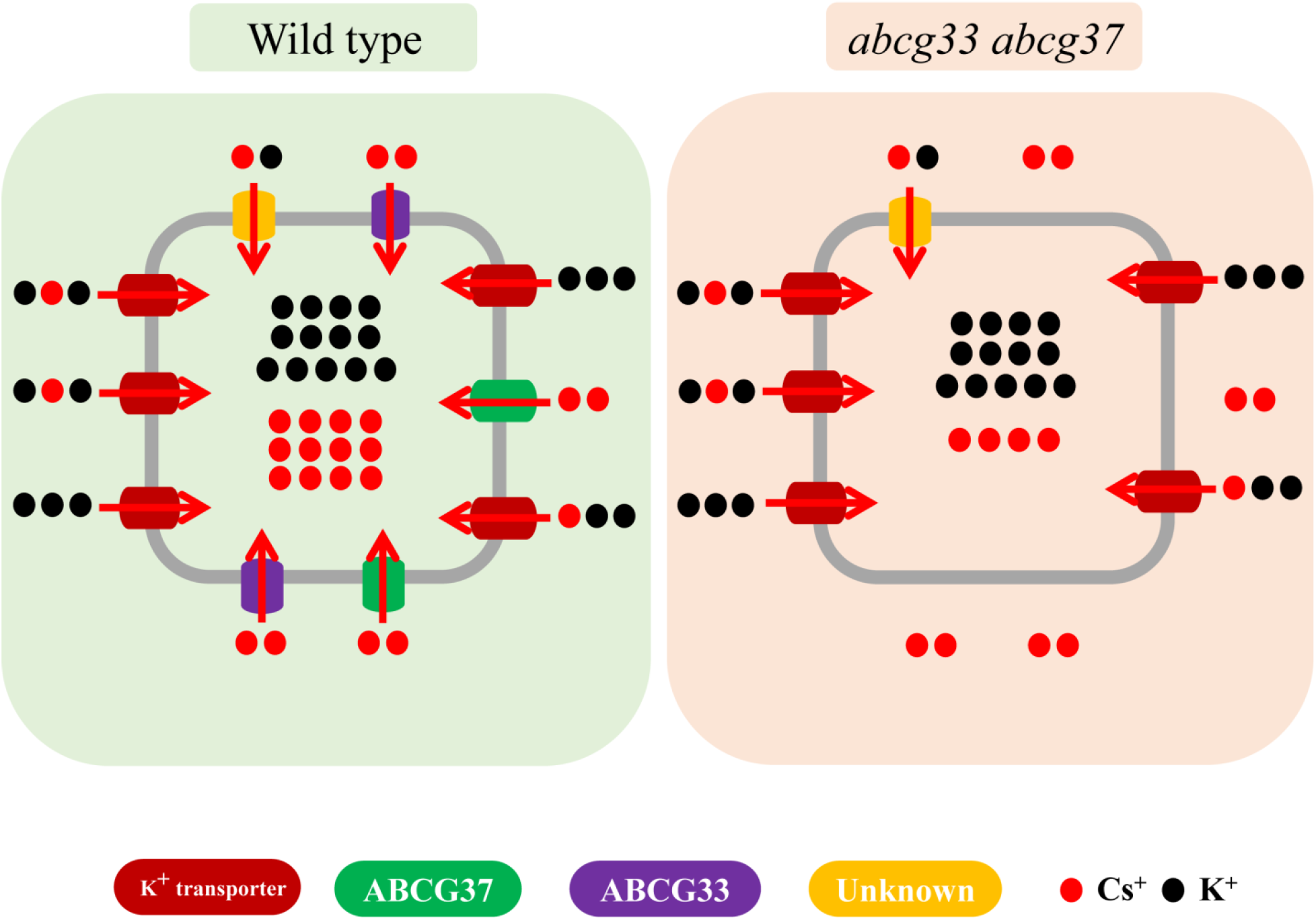
Graphical model of cesium uptake. Schematic representation of current working model of Cs^+^ uptake in Arabidopsis. Left panel demonstrates the typical Cs^+^ uptake inside wild-type (Col-0) cell using various transporters, including ABC proteins. Right panel shows the loss of ABCG33 and ABCG37 in double knockout mutant resulting in reduced Cs^+^ uptake. K^+^ transporters are presented in red, ABCG33 in purple, ABCG37 in green and yet unidentified Cs^+^ transporters are in yellow.

## Methods

### Plant materials

All lines are in the Columbia background of *Arabidopsis thaliana*. *abcb1*/*mdr1*, *abcb2*/*mdr2*, *abcb3*/*mdr3*, *abcb4*/*mdr4*, *abcb5*/*mdr5*, *abcb6*/*mdr5*, *abcb6*/*mdr6*, *abcb7*/*mdr7*, *abcb9*/*mdr9*, *abcb10*/*mdr10*, *abcb11*/*mdr8*, *abcb12*/*mdr16*, *abcb13*/*mdr15*, *abcb14*/*mdr12*, *abcb15*/*mdr13*, *abcb16*/*mdr18*, *abcb17*/*mdr19*, *abcb18*/*mdr20*, *abcb19*/*mdr11*, *abcb20*/*mdr14*, *abcb22*/*mdr21* were provided by Edgar Spalding, Department of Botany, University of Wisconsin-Madison and *abcg37-1*/*pdr9-1* was provided by Willian M. Gray, Department of Plant Biology, University of Minnesota. Other T-DNA insertion mutants were obtained from ABRC (Arabidopsis Biological Resource Center), and listed in the Supplemental Table 2.

ABCG37-GFP (Růžička et al., 2010), ABCG33-GFP (Schuetz et al., 2014), PIN2-GFP (Xu and Scheres, 2005), EGFP-LTI6b (Kurup et al., 2005) and *pCyclinB1;1-GUS* were described elsewhere (Colón-Carmona et al., 1999).

Three independent double knockout mutants (*abgc37-2 abcg33-1*) were generated by crossing and the homozygous lines were selected through genotyping. Primers used for genotyping are listed in the Supplemental Table 1.

### Growth conditions

Surface-sterilized seeds were placed in round, 9-cm Petri plates on modified Hoagland medium (Baskin and Wilson, 1997) containing 1% (w/v) Sucrose and 0.8% (w/v) agar (Difco Bacto agar; BD Laboratories, http://www.bd.com). Two days after stratification at 4°C in the dark, plates were transferred to a growth chamber (NK System; LH-70CCFL-CT, http://www.nihonika.co.jp) at 23°C under continuous white light (at an irradiance of ∼100µmol m^-2^s^-1^). The seedlings were grown vertically for 3 days. For cesium chloride treatment, 3-days-old seedlings were transferred to cesium chloride containing plates of various concentrations and incubate for various time points. Root elongation data represent the root elongation after transferring to the corresponding media and mentioned concentrations. To measure the root elongation, photographs of plates were taken using a digital camera (Power Shot A640, Canon, http://canon.jp) and analyzed by an image analyzing software ImageJ (http://rsb.info.nih.gov/ij/).

For auxin response assay, 4-d-old light grown Col-0, *abcg37-1*, *abcg37-2*, *acbg33-1* and *abcg37-2 abcg33-1* seedlings were transferred to DMSO, 15nM and 30nM IAA containing plates and kept at 23°C under yellow light for three days. After the incubation, root elongation was measured as described above.

### Chemicals

Cesium chloride was purchased from Sigma Chemicals (www.sigmaaldrich.com). Propidium idodide was purchased from Invitrogen (www.thermofisher.com). The carrier-free ^137^Cs solution (3.7MBq mL^-1^) was purchased from Eckert & Ziegler (https://www.ezag.com). Other chemicals were purchased from Wako, Japan (www.wako-chem.co.jp).

### Kinematic analysis

Kinematic analysis was performed as described earlier (Rahman et al., 2007). In brief, seedlings were grown vertically for 4 days after stratification. On day 4, seedlings were transferred to plates supplemented with or without and 1.5mM cesium chloride and grown vertically for another 3 days. Root elongation was measured by scoring the position of the root tip on the back of the Petri plate once per day. After the end of the incubation, cortical cell length was measured using light microscope (Nikon Diaphot) equipped with a digital camera control unit (Digital Sight [DS-L2]; Nikon, Japan, https://www.nikon.co.jp). To ensure newly matured cells were scored, cells were measured for the root zone, where root hair length was roughly half maximal. The length of 10 mature cortical cells was measured from each root, with 8 roots used per treatment. The cell production rate was calculated by taking the ratio of root elongation rate and average cell length and average cell length for each individual and averaging over all roots in the treatment. The data were obtained from at least three independent biological replicates.

### GUS staining

GUS staining was performed as described earlier (Okamoto et al., 2008). In brief, 3-d-old seedlings were transferred to 1.5mM cesium chloride containing agar plate and grown vertically at 23°C under continuous white light. After three days incubation, seedlings were transferred to GUS staining buffer (100 mM sodium phosphate, pH 7.0, 10mM EDTA, 0.5mM potassium ferricyanide, 0.5 mM potassium ferrocyanide, and 0.1% Triton X-100) containing 1 mM X-gluc and incubated at 37°C in the dark for 3 h. The roots were imaged with a light microscope (Nikon Diaphot) equipped with a digital camera control unit (Digital Sight [DS-L2]; Nikon, Japan).

### Propidium Iodide staining

0.01mg/ml Propidium Iodide (PI) solution was used to stain the root and subjected to confocal microscope imaging immediately. Roots were mounted in PI solution containing coverslip and imaged using a Nikon laser scanning microscope (Eclipse Ti equipped with Nikon C2 Si laser scanning unit, https://www.nikon.co.jp) with a X20 objective. The imaging was performed within 5 minutes from the start of the PI staining. The experiment was repeated three times.

### Gene expression analysis

3-d-old vertically grown *Arabidopsis thaliana* seedlings were transferred to 1.5 mM cesium chloride containing agar plate and incubated at 23°C for additional three days. After the treatment, RNA was extracted from the root tissue using RNA Extraction Kit (APRO Science, Japan, www.aprosci.com) with on-column DNA digestion to remove residual genomic DNA using RNase-free DNase according to manufacturer’s protocol. Extracted RNA was tested for quality and quantity. Each RNA concentration was normalized with RNase free water. 500 ng RNA was used to synthesize cDNA using Rever Tra Ace qPCR RT master mix (Toyobo, Japan, www.toyobo-global.com). Quantitative PCR reactions were performed using the Takara TP-850 thermal cycler (Takara Bio, Japan, www.takara-bio.com) and SsoAdvanced^TM^ Universal SYBR^®^ Green Supermix (BIO-RAD, USA, www.bio-rad.com.The reaction was performed as per manufacturer’s instruction. For quantification of *ABCG* expression, we used the 2-^ΔΔ^CT (cycle threshold) method with a normalization to the ef1α expression (Hanzawa et al., 2013). Data were obtained from three biological replicates. Primers used for the gene expression analysis are listed in Supplemental Table 1.

For RT-PCR analysis (Supplemental Figure 4), RNA was extracted from 7-day-old vertically grown seedlings and cDNA was prepared as described above. RT-PCR was performed using GoTaq^®^ DNA polymerase (Promega, https://www.promega.com) and corresponding primers for 28 cycles. Primers used for the gene expression analysis are listed in the Supplemental Table 1.

### Live-Cell imaging

To image GFP, the 3-day-old seedlings were transferred to 1.5mM cesium chloride containing plates and incubated at 23°C under continuous light for three days. Images were taken in every 24h time interval up to 72h (Figures 4A and 4B; Supplemental Figure 8). For colocalization study, five-day-old ABCG37 and ABCG33-GFP seedlings were grown and stained with 0.01mg/ml propidium iodide (Supplemental Figure 8). After mounting on a large cover glass, the roots were imaged using a Nikon laser scanning microscope (Eclipse Ti equipped with Nikon C2 Si laser scanning unit) with a X20 objective. Same confocal settings were used for each group of experiments. Fluorescence intensities were measured by drawing same region of interest (ROI) in the images obtained from live-cell imaging using Image J software.

### Cesium transport assay

4-day-old light grown seedlings were incubated in 0.1µM, 10µM, 1.5mM and 15mM ^137^Cs^+^ containing liquid Hoagland solution for 2h both in presence (6mM K^+^) or absence of potassium, transferred over Nylon mesh and briefly washed with liquid Hoagland solution three times. 10mm root tip from 10 seedlings were collected for each sample and placed into the scintillation vial with 500µL MicroScint 40 (Perkin Elmer, Inc., USA, www.perkinelmer.com). The uptake of ^137^Cs^+^ was determined with a NaI scintillation counter (ARC-300, Aloka, Japan, www.hitachi-aloka.co.jp). The experiment was repeated for three times.

### Measurements of ion contents

Three-day-old Col-0and double knockout mutant seedlings were transferred to control and 1.5mM cesium chloride containing plates with variable K^+^ concentrations, and incubated for three days at 23°C under continuous white light. Whole seedlings were collected, washed three times with ultrapure water, soaked in the paper towel and the fresh weight was measured. Samples were dried at 65°C and digested using ultrapure nitric acid (Kanto Chemical, Japan, https://www.kanto.co.jp) at 95°C for 600 min. Ion contents were determined using an inductive coupled plasma mass spectrometer (ICP-MS, NexION 350S, Perkin Elmer, Japan).

### Cesium transport assay in *Saccharomyces cerevisiae*

To evaluate the Cs^+^ transport by ABCG37, we performed the yeast culture experiment under various Cs^+^ concentrations. The ABCG37 cDNA was cloned into the pENTR/D-TOPO vector (Invitrogen, USA). Primers (ABCG37_F, ABCG37_R) used for cloning are listed in the Supplementary table 1. Subsequently, ABCG37 cDNA was transferred into the destination vector pYES-DEST52 (Invitrogen, USA) to construct the expression vector for yeast transformation. The yeast strain, INVSc1 (MATa his3D1 leu2 trp1-289 ura3-52 MAT his3D1 leu2 trp1-289 ura3-52), was transformed with the pYES-DEST52-ABCG37. After confirming the growth in the overnight culture with 2%, SC-URA medium, the yeast strains were washed with RO water twice, then cultured in SC-URA medium with 2% galactose in 0.1µM, 100 µM and 1.5mM of Cs^+^. After 24 hours, OD600 of each sample was measured to adjust equal number of cells for each treatment. The collected samples were washed with RO water twice and digested by 13N nitric acid. Cs^+^ content in the sample solution was measured by inductively coupled plasma mass spectrometry ICP-MS (NexION 350S, PerkinElmer).

### Bioinformatics analysis

Mutational information of ABCG37 mutants were collected from SALK T-DNA repository (http://signal.salk.edu/cgi-bin/tdnaexpress) and schematic diagram of mutants were drawn based on Exon-Intron Graphic maker (http://wormweb.org/exonintron).

Protein sequences of ABCG transporters were collected from TAIR database (https://www.arabidopsis.org/index.jsp) and phylogenetic tree was constructed using online Clustal Omega (https://www.ebi.ac.uk/Tools/msa/clustalo/) tool.

## Acknowledgements

The authors thank Edgar Spalding (Department of Botany, University of Wisconsin – Madison), William M. Gray (college of biological sciences, University of Minnesota), Angus Murphy (University of Maryland, USA), Lacey Samuels (University of British Columbia, Canada), and Ben Scheres (University of Utrecht, The Netherlands) for sharing materials. This work was partially funded by Iwate University President Fund (A.R.), UGAS, IU Research fund 2017 (M.A.A.), and Japan Science and Technology Agency (JST) [PRESTO # 15665950] (K. T.). M.A.A. was supported by MEXT fellowship.

## Author Contribution

A.R. designed and supervised the experiments. Initial screening of ABCB and ABCG mutants was done by S.K. M.A.A. and K.I. performed the experiments. R.S. and K.T. assisted for the Cs^+^ uptake assay, ICP-MS experiment and data analysis. A.R. and M.A.A. and K.T. analyzed the data. A.R. and M.A.A. wrote the manuscript.

